# Residue-specific insights into (2x)72 kDa tryptophan synthase obtained from fast-MAS ^1^H-detected solid-state NMR

**DOI:** 10.1101/2021.05.12.443859

**Authors:** Alexander Klein, Petra Rovó, Varun V. Sakhrani, Yangyang Wang, Jacob Holmes, Viktoriia Liu, Patricia Skowronek, Laura Kukuk, Suresh K. Vasa, Peter Güntert, Leonard J. Mueller, Rasmus Linser

## Abstract

Solid-state NMR has emerged as a potent technique in structural biology, suitable for the study of fibrillar, micro-crystalline, and membrane proteins. Recent developments in fast-magic-angle-spinning and proton-detected methods have enabled detailed insights into structure and dynamics, but molecular-weight limitations for the asymmetric part of target proteins have remained at ~30-40 kDa. Here we employ solid-state NMR for atom-specific characterization of the 72 kDa (asymmetric unit) microcrystalline protein tryptophan synthase, an important target in pharmacology and biotechnology, chemical-shift assignments of which we obtain *via* higher-dimensionality, 4D and 5D solid-state NMR experiments. The assignments for the first time provide comprehensive data for assessment of side chain chemical properties involved in the catalytic turnover, and, in conjunction with first-principles calculations, precise determination of thermodynamic and kinetic parameters is demonstrated for the essential acid-base catalytic residue βK87. The insights provided by this study expand by nearly a factor of two the size limitations widely accepted for NMR today, demonstrating the applicability of solid-state NMR to systems that have been thought to be out of reach due to their complexity.

## Introduction

Complementing the insights from crystallography, optical spectroscopy, and simulation, NMR spectroscopy has been an invaluable asset for mechanistic studies of enzymatic catalysis. In addition to atomic-resolution access to protein dynamics and domain motion readily available for resonance-assigned targets, the individual chemical shifts themselves are a prime source of information reflecting chemical properties, which are often poorly represented in structural data obtained from other techniques. The site-specific assignment of chemical shifts, however, has remained the main bottleneck for any downstream NMR investigation for most of the potential targets. Proteins much larger than 30 kDa are difficult to assign specifically by solution state NMR due to slow tumbling, which results in broadened resonances and low intensity upon INEPT-based magnetization transfer. The latter is a particular problem for more elaborate strategies, like higher-dimensionality experiments, which are needed for large proteins to attain sufficient resolution for unambiguous sequential correlation but require a multitude of magnetization transfers. Although site-specific and methyl labelling schemes^[1,2]^ are useful to answer specific biological questions for larger systems, the usual repertoire of multifaceted NMR-spectroscopic experiments becomes ineffective.

Over the last several years, ^1^H-detected solid-state NMR has been established as a robust technique for providing insights into protein structure, interactions, and dynamics in systems of increasing complexity.^[3]^ Innovations in sample preparation, most notably deuteration, ^1^H back-exchange, and paramagnetic doping^[4–8]^, in combination with hardware enabling very high magic angle spinning (MAS) frequencies and smart spectroscopic approaches, have facilitated access to atom-specific chemical shifts in increasingly challenging target proteins.^[3,9–11]^ Whereas limited signal intensity can often be compensated by extended measurement times and high magnetic fields, the devastatingly high probability of resonance overlap and the resulting ambiguities in the assignment of proteins with molecular weight exceeding 40 kDa have, again, seemed an insurmountable problem. In the solid state, in contrast to solution NMR of globular proteins, however, transfer efficiencies are independent of molecular weight. Increasingly complex solid-state NMR approaches like 4D experiments, particularly in combination with proton-detected methodology, have recently been demonstrated to reduce redundancy in experiments for spectral assignments^[9,12–17]^, structure calculation^[18–23]^, and characterization of protein dynamics^[13]^. For systems containing individual asymmetric units with molecular weights of 40 kDa and above, however, even these experiments tend to show significant ambiguity, and backbone assignments for such targets are almost completely absent. Most proteins of medical, biological, or biotechnological interest are far more complex than those studied to date, making effective methodological developments to expand the accessible size range of NMR spectroscopy potentially one of the most profitable and impactful tasks in structural biology. Here we demonstrate that the use of proton-detected, fast-magic-angle-spinning solid-state NMR, in particular of higher dimensionality (>3D), makes it feasible to disambiguate sequential shift correlations for partial resonance assignments, gives access to relaxation data, and allows a direct use of experimental data for mechanistic studies even for very high protein molecular weights.

Fig. 1A depicts the crystal structure of the *S. typhimurium* tryptophan synthase (TS) complex, a pyridoxal 5’ phosphate (PLP)-dependent enzyme. TS is an αββα heterodimer with an αβ asymmetric unit of 72 kDa or 665 amino acids (total molecular weight 144 kDa).^[24]^ The family of PLP-dependent enzymes catalyzes a wide variety of chemical transformations including transamination, racemization, decarboxylation, elimination, and substitution, causing a widespread interest in deciphering the details of the catalytic cycle.^[25–29]^ The large number of PLP enzymes and their crucial metabolic functions make them significant drug targets for the treatment of diseases including tuberculosis, epilepsy, and Parkinson’s disease.^[30–32]^ TS itself is both an important drug target in the context of continuously emerging bacterial antibiotics resistance^[33]^ and of great interest in biotechnology^[34]^ as an enantiospecific source of a large variety of unnatural amino acids and their derivatives.^[35–37]^ Wildtype TS catalyzes the final two steps in tryptophan biosynthesis: production of indole from IGP at the α site, and its subsequent condensation reaction with L-serine to give L-tryptophan^[24,38–41]^ (Fig. 1B). For TS, X-ray structural data is abundant. But the rational design of therapeutic agents and the understanding and engineering of TS catalysis, in particular regarding the β-subunit enzymatic reaction, hinges on the availability of detailed knowledge of the chemical and electrostatic properties of the active site. The latter are strongly influenced by the protonation, hybridization, and tautomeric states of the active-site side chains and substrates, which are not directly determined by protein crystallography but are accessible from NMR chemical shifts. In particular, the chemical properties of the βK87 Schiff base have remained an important focus with implications for the reaction intermediates in this and other PLP-dependent enzymes.[40],[41]

**Fig. 1:**
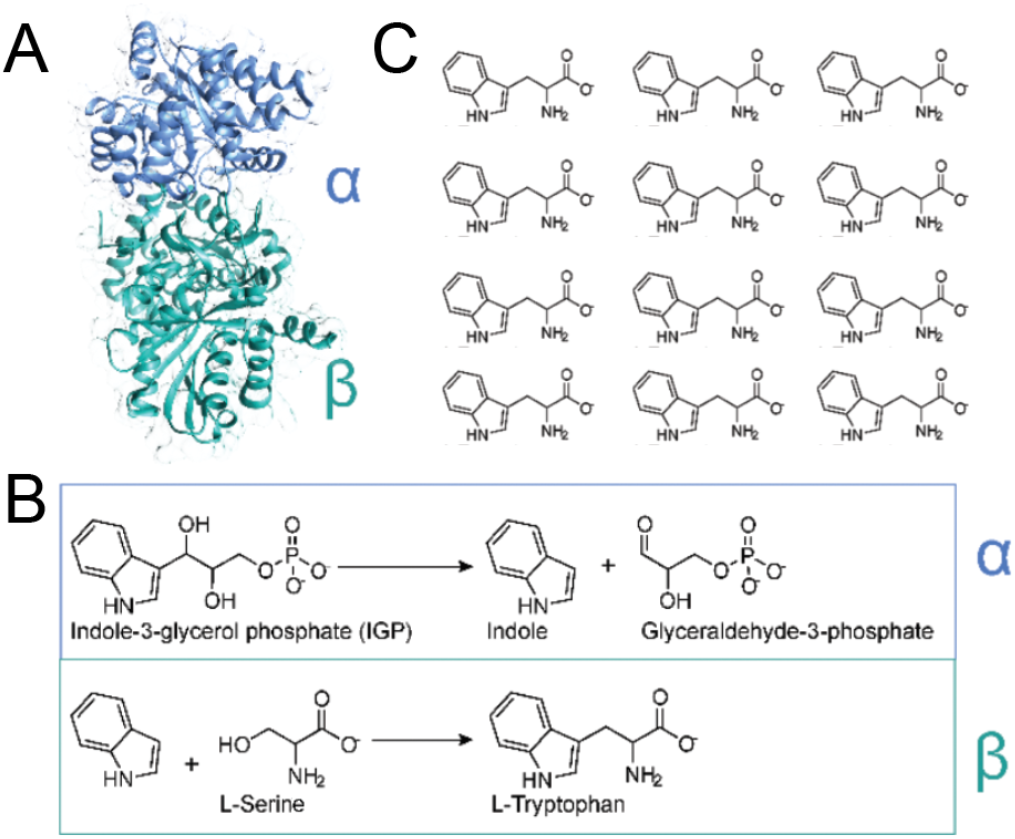
Tryptophan synthase (TS) is an αββα heterodimer with an asymmetric unit of 72 kDa. **A)** Topology of the αβ asymmetric unit for the internal aldimine resting state (pdb 4ht3).^[24]^ **B)** The reactions catalyzed by wildtype TS in the α and β subunits that comprise the final two steps of tryptophan biosynthesis. **C)** A selection of substrates and products of engineered TS enzymes for potential use in biotechnological applications. The lower right compound represents thaxtomin A, a natural product synthesized from 4-nitroTrp (lower left)^[35,36,43]^.

TS represents an enzyme 1.7x the size of the current record of high-molecular-weight proteins in solid-state NMR^[42]^ (just considering the 72 kDa *asymmetric* portion of the enzyme), such that protein chemical shifts have been remaining effectively absent, and NMR studies on TS have been limited to selectively labeled substrates and cofactors of the enzyme.^[39–41]^ Providing partial but substantial assignments, here we show that state-of-the-art, proton-detected, fast-magic-angle-spinning solid-state NMR allows studies of catalytic details that so far have been deemed inaccessible with current methodology.

## Results

Successful NMR assignments in proteins of increasing molecular weight hinges on sufficient dispersion of resonances. Higher-dimensionality (>3D) experiments are of limited applicability, however, due to their large number of transfer steps and indirect evolution times. Compared to solution NMR, solid-state NMR ameliorates this bottleneck, as cross polarization leverages effective magnetization transfer within short times and without the refocusing periods needed for INEPT transfers and, during indirect evolution times, *T*_2_ relaxation times being comparably long in case fast MAS and/or high levels of deuteration are used. It has far-reaching consequences that the framework of solid-state NMR is fundamentally independent of molecular weight, allowing experiments to be applied with the same overall efficiency to proteins of any size and thus increasingly outperform solution NMR experiments for higher molecular-weight and experiment complexity/dimensionality^[9,44]^.

To tackle *S. typhomurium* tryptophane synthase, we used a single, triply ^2^H, ^13^C, ^15^N-labeled and micro-crystalline sample in a 1.3 mm rotor at 700 MHz proton Larmor frequency proton-back-exchanged over the course of multiple weeks, and subjected it to a suite of established as well as newly engineered higher-dimensionality (4D and 5D) solid-state NMR experiments. (In particular, a 4D hCACONH, 4D hCOCANH, 4D hCACBcaNH, 4D hCACBcacoNH, 5D HNcoCANH, a 4D hCCNH (side chain to backbone correlation), and a pseudo-4D ^15^N *R*_1ρ_-edited hCONH series, see preparative and experimental details, as well as individual relaxation decays in the SI.) Fig. S1 shows a set of spectral excerpts for K355 as an exemplary residue. Even though utilization of fully protonated samples (at >100 kHz) would be the desired labeling scheme to avoid incomplete back-exchange, that labelling unfortunately turned out to provide sensitivities that would not have allowed acquisition of the experiments within a manageable timeframe. (See the SI for a specific comparison between 1.3 and 0.7 mm samples.) Assignments were supported by state-of-the-art computational capability via FLYA^[45]^, using a suitably modified set of parameters within the existing software environment. This computation-aided strategy enables a residue-specific, quantitative assessment of assignment quality in an iterative way, which becomes indispensable for the complexity of experimental data obtained and the large number of residues to converge with high reliability. As in particular 5D spectra have not been utilized in solid-state NMR (apart from APSY approaches),^[9]^ the SI contains a brief description of setting the experiments up and processing the data acquired.

Just like manual assignment but with the possibility to validate combinations of possible assignment combinations faster, FLYA-aided assignment benefits from the high redundancy introduced by the combination of multiple higher-dimensionality approaches. Fig. 2 depicts the reliably obtained assignments of TS. (See the SI for a list of the chemical shifts obtained.) Counting only high-confidence, “strong” assignments, at least 369 *completely* backbone*-* (H^N^, N^H^, CO, Cα and Cβ-assigned residues could be assigned, which corresponds to 59% of the non-proline residues. (Noteworthily, this refers to very conservative validation with stringent exclusion criteria in FLYA with respect to next-neighbor assignments and only when at least all of H^N^, N, Cα, CO, and Cβ resonances within these residues are marked “strong”). Additional 121 residues are assigned where part of these individual resonances within a residue are unambiguous, but not all of the five nuclei above reach the “strong” attribute. Thus in total, 491 residues are obtained in which at least part of these nuclei bear a “strong” assignment in FLYA. Although only representing a partial assignment, this reasonably large number (considering not only the excessively high molecular weight and associated limitations discussed but also the highly dynamic enzymatic machinery and expected exchange broadening in areas like the COMM domain and the αL2 and αL6 regions) is an unprecedented basis for gathering site-specific information on chemical properties in TS. Fig. 2A also shows the secondary-structure elements derived by TALOS-N^[46]^ based on the assigned chemical shifts compared to those from the crystal structure (PDB: 4ht3^[24]^). Secondary chemical shifts are shown in Fig. S11. Without further treatment, NMR chemical shifts are direct reporters on the chemical and electrostatic properties of individual sites within the protein, including hybridization, tautomerization, and H-bonding states. Chemical shifts in TS are of major interest for gathering details of the prototypical PLP-dependent catalytic reaction, but have been available to date only for specifically labeled cofactor, substrates, and analogues for specific complexes and inhibited reaction intermediates. Chemical shift data represent the foundation for investigating the catalytic mechanism, often with assistance from crystallography and DFT calculations.^[24,40,47,48]^ For the first time, this study can provide chemical shifts of backbone and sidechain atoms of many of the residues within the protein that are directly involved in the catalytic mechanism. Such residues are located within the active sites of the two subunits as well as in the interface between the subunits, through which allosteric communication between the active sites occurs. For example, G500 (βG232) in the β-subunit active site has a strongly downfield shifted H^N^ chemical shift of 10.58 ppm, evincing its hydrogen bond to the cofactor phosphate group. Similarly, E618 (βE350) shows a remarkable H^N^ chemical shift of 10.97 ppm, which reflects the salt bridge it can form to K650 (βK382) in conjunction with a hydrogen bond between its H^N^ and its own sidechain.

**Fig. 2:**
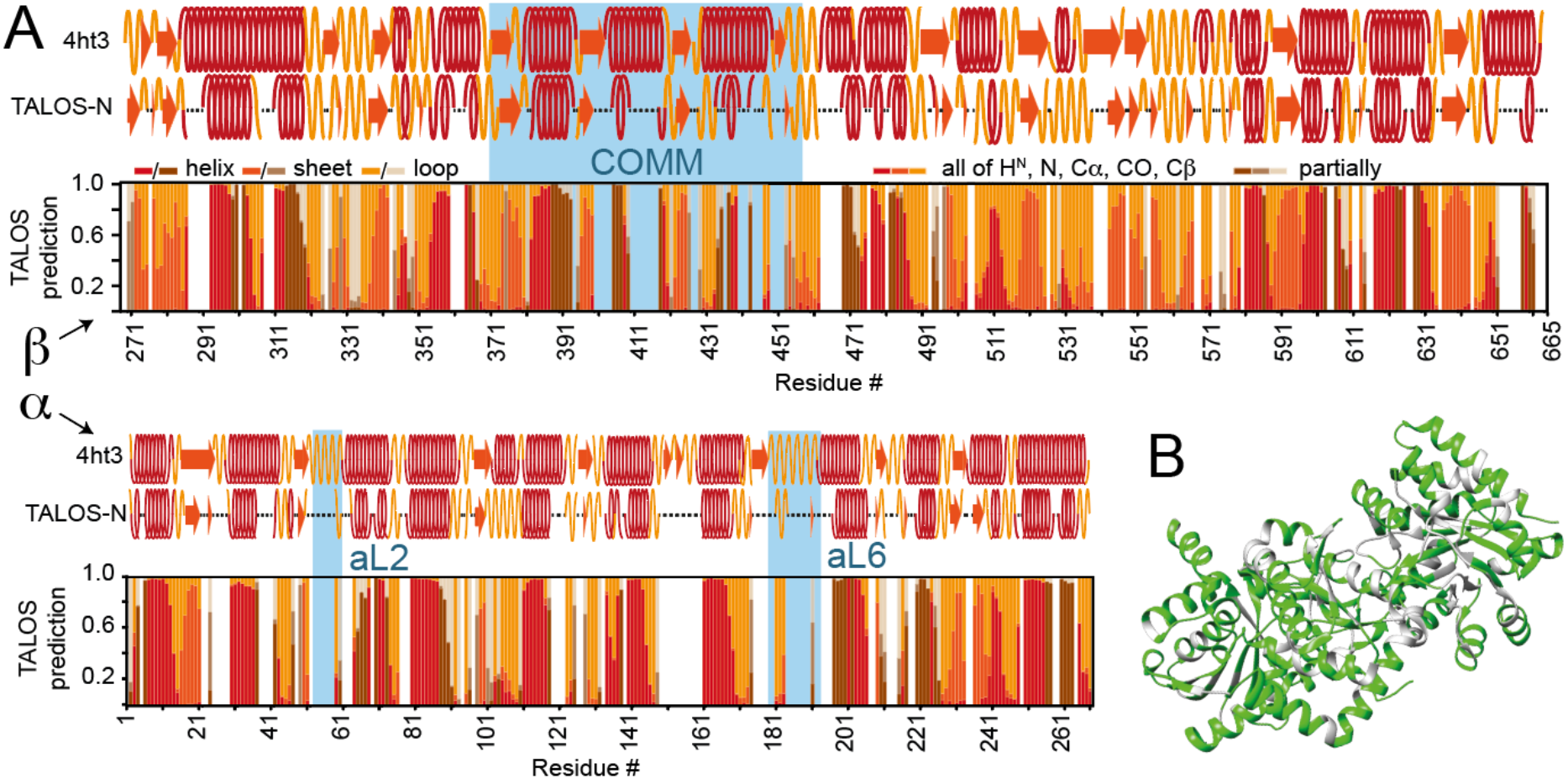
Chemical-shift assignment in TS. **A)** Assigned residues for the β-(top) and α-subunit (bottom) and analysis with respect to the secondary structure predicted by TALOS-N, shown only for all confident assignments. Red-tone colors denote high-confidence assignment in FLYA quality assessment, but only in case all of H^N^, N, Cα, CO, and Cβ are assigned as “strong”, “strong” being defined very conservatively (see the SI). Brown colors denote that at least part of the mentioned five nuclei have “strong” assignment. The TALOS results (red: helix; dark orange: sheet; orange: loop) are compared with the secondary structure elements found in the crystals structure 4ht3[24] (top row). Highlighted in light blue are domains considered to undergo major conformational changes upon ligand binding at different states of the catalytic cycle[24]. Short stretches, e. g., 501 to 511 or 529 to 533 show minor differences between TALOS and the actual crystal structure, while SCS (Fig. S11) are in agreement. These deviations are thought to arise from TALOS prediction known to be improper in case of short stretches or at the edges of assigned regions. **B)** Residues with strong assignments for all of the above-mentioned nuclei marked on the crystal structure (green, pdb: 4ht3).

The catalytic pocket of the β-subunit has been of extensive scientific interest for understanding overarching properties of PLP-based enzymes as well as of TS specifically, [25–29] particularly to foster engineered variants transforming a broader range of substrates.[35–37] Fig. 3A depicts the interesting residues and H-bond network around the cofactor at the beginning of the reaction (see Fig. 3B), at the same time color-coded by their ^15^N *R*_1ρ_ rates. The active-site chemical shifts are particularly interesting in combination with first-principles calculations when comparing different catalytic states or mutations.^[24,40,47,48]^. For example, K355 (βK87) in the β-subunit active site, which holds the cofactor through a Schiff base linkage, shows a noticeably unique Cδ chemical shift of 31.7 ppm. Likewise, the adjacent H354 (βH77) shows a Cβ chemical shift of 30.7 ppm, which suggests an elevated pK_A_ of the imidazole sidechain in this environment. S645 (βS377) on the other side of the β-subunit active site, which is sandwiched by two short H-bonds (Fig. 3) to the PLP cofactor and S619 (βS351), has a noteworthy Cαchemical shift of 62.5 ppm. Simultaneously being an H-bond acceptor for S619, the S645 (βS377) Cβ shift amounts to (only) 62.7 ppm. S619 (βS351) has rather usual Cα/Cβ shifts of 59.8/62.1 ppm, even though the inter-Ser H-bond has a shorter donor-acceptor distance.

**Fig. 3:**
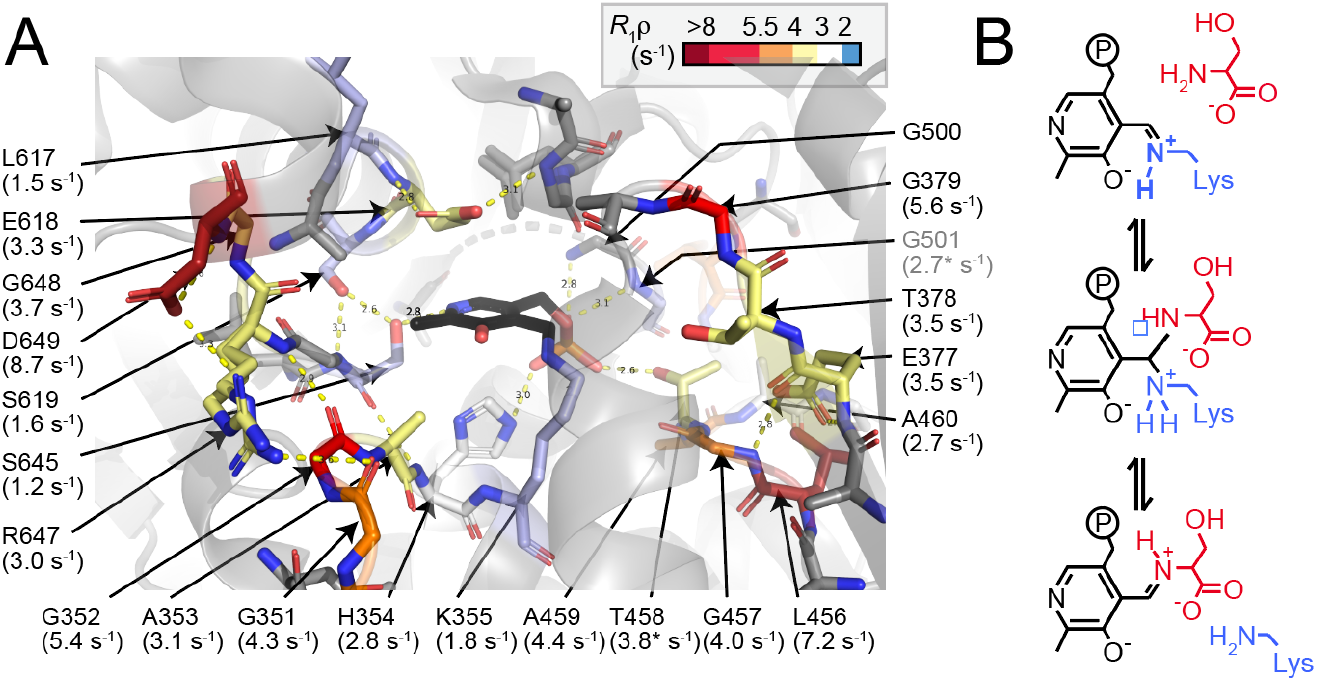
Access to chemical shift information specific to β-subunit catalysis. **A)** Assigned residues in the active site of the β-subunit, with ^15^N *R*_1ρ_ rates annotated and color-coded on the X-ray structure (pdb 4hpj). The color code is shown on the top. Residues in gray sticks are assigned but not sufficiently resolved in the pseudo-4D relaxation experiments. Non-assigned residues are shown without sticks. The PLP cofactor is shown in black. H-bonds (yellow dashes) are shown with their inter-heteronuclear distances (in Å). **B)** Initial step of the catalytic cycle, with the protonation state of the Schiff base (top) of interest.

Most importantly, all shifts for the lysine sidechain covalently bonded to the PLP cofactor are now available, giving direct experimental access to the question of the linking Schiff base equilibrium protonation state and the associated energies that characterize the initial step of the catalytic cycle.^[41]^ Protonation of the Schiff base has been considered as a means to activate the C4’ carbon of the cofactor for nucleophilic attack by serine (Fig. 3B).^[40,49–51]^ Whereas the ^15^N ζ shift has been addressed in isolation previously through selective labeling and heteronuclear-detected low-temperature ssNMR,^[39]^ no carbon sidechain or proton shifts were obtained. By contrast, the proton-detected 4D and 5D sequential backbone experiments here afford K355 (βK87) backbone shifts, enabling side chain carbon assignments from Cα through Cɛ via the 4D hCCNH, which link to N and the H ζ proton in the H/N via a long-range H/C correlation (Fig. 4A).

**Fig. 4:**
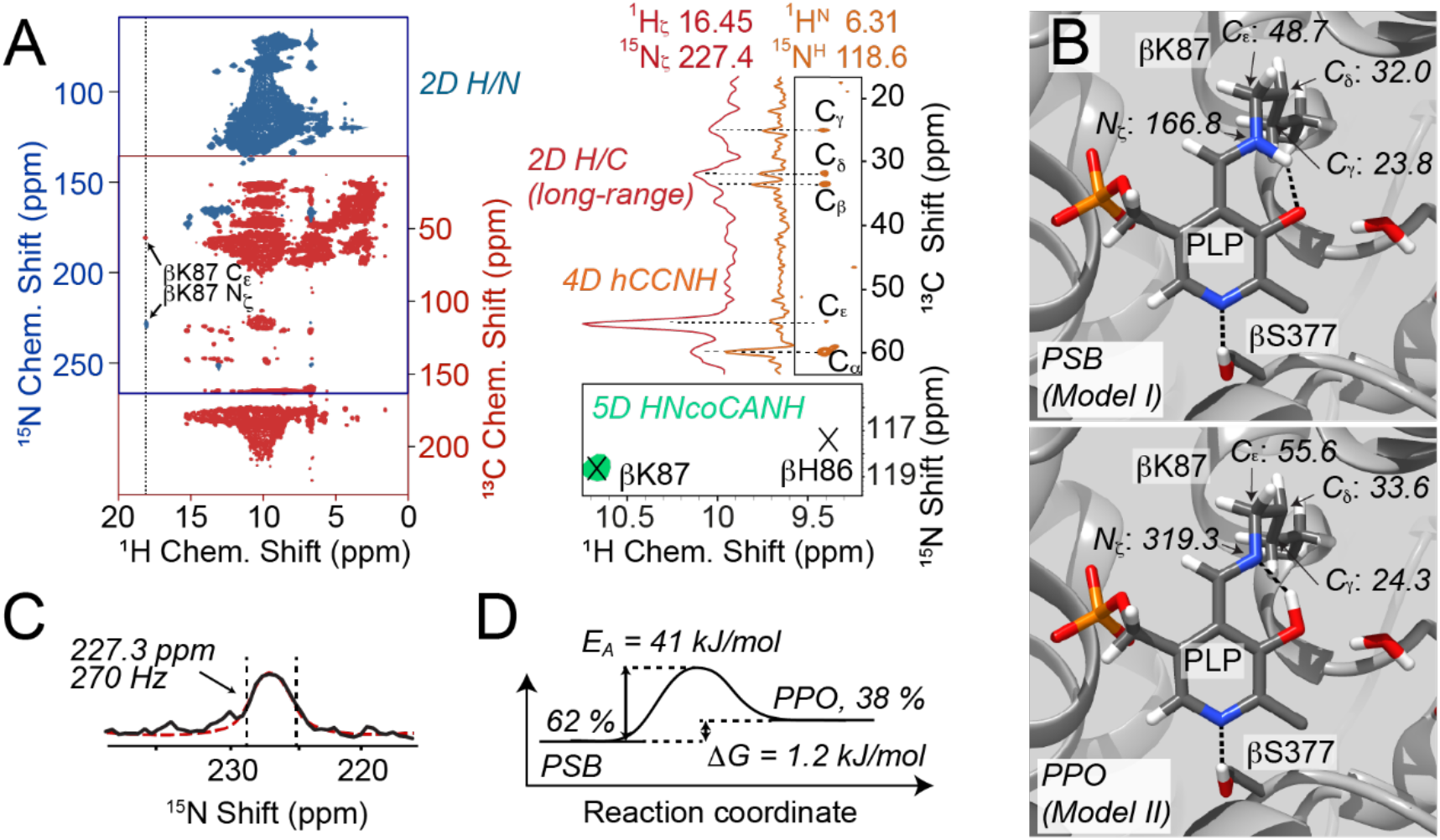
Analysis of the tautomeric equilibrium at the βK87 Schiff base in the active site of the β-subunit. **A)** Complete assignment of βK87 via backbone assignments (5D HNcoCANH, green, representing the backbone experiments), sidechain carbon assignments (4D hCCNH, yellow), H/N correlation (blue), and a connecting 2D long-range H/C correlation (red). **B)** Shifts from *ab-initio* calculations for the active site with protonated Schiff base (PSB, model I, top) and protonated phenolic oxygen (PPO, model II, bottom). **C)** Linewidth analysis for the ^15^N of the Schiff base (spectrum with 50 Hz exponential decay in black, Lorenzian in red,). **D)** Resulting free-energy profile of the tautomeric exchange of the Schiff base at 30 °C. The plane from the 5D HNcoCANH depicted in A) is extracted at H,N,Cα shifts 9.44/117.1/58.9, with the ^1^H coordinates shown folded in from 6.3 ppm (indirect H^N^).

This series of active-site shifts confer unprecedented experimental insights. The intermediate value of Nζ found here at 227.3 ppm at 30°C compared to the previously reported experimental value of 202.3 ppm at -10°C^[40]^ suggests a dynamic tautomeric exchange between protonated and deprotonated Schiff base forms. Quantitatively, the set of experimental chemical shifts can now be used as restraints to evaluate precise computational models of structure and dynamics in the active site using “NMR-assisted crystallography,” the combined application of solid-state NMR, X-ray crystallography, and first-principles computational chemistry^[40,47,52–55]^. As detailed in the SI, this synergistic approach uniquely points to a fast-exchange model involving the protonated Schiff-base (PSB) and protonated phenolic oxygen (PPO) tautomeric forms (Fig. 4B), quantitatively confirming this exchange and uniquely identifying the exchange partner. Table 1 summarizes select experimental and first-principles predicted shifts for these two species. Individually, the PSB (model I) and PPO (model II) tautomers show little congruency with the experimental data as quantified using the reduced-χ^2^ statistic^[40]^, while the fast-exchange equilibrium between them (model III), with a reduced-χ^2^ of 0.5, is well in line with the experimental measurements.

**Table 1:** Experimental and first-principles predicted chemical shifts (ppm) for the PSB (Model I), PPO (Model II), and their best-fit two-site exchange (Model III).

|  | Experimental | Model I | Model II | Model III |
| --- | --- | --- | --- | --- |
| K355 (βK87) Nζ / SBN | 227.3 | 166.8 | 319.3 | 224.6 |
| K355 (βK87) Cγ | 24.9 | 23.8 | 24.3 | 24.0 |
| K355 (βK87) Cδ | 31.7 | 32.0 | 33.6 | 32.6 |
| K355 (βK87) Cε | 52.3 | 48.7 | 55.6 | 51.3 |
| PLP N1 | 294.7 | 304.4 | 302.7 | 303.8 |
| PLP C2 | 159.6 | 160.0 | 151.3 | 156.7 |
| PLP C2' | 20.4 | 21.3 | 20.3 | 20.9 |
| PLP C3 | 168.3 | 171.9 | 156.8 | 166.2 |
| Reduced $\chi^2$ | - | 26.9 | 69.7 | 0.50 |

From these calculations, the equilibrium populations for the PSB and PPO forms directly arise as 62% and 38%, respectively, indicating a free-energy difference of only 1.2 kJ/mol between the tautomers at 30°C. The corresponding populations at -10°C are 78% and 22%. The temperature dependence of the populations is inconsistent with a simple two-state model for the exchange, unless an excessive entropy difference is introduced. We hypothesize that this shift in populations is induced in part by large-scale protein dynamics between the open and closed conformations of the β-subunit. The open conformation is necessary for the free diffusion of substrate into the active site, but also establishes an aqueous environment proximal to the cofactor that favors the Zwitterionic, PSB form. Closed conformations largely exclude water from the active site, favoring the neutral Schiff base, PPO form. It is expected that open and closed forms of the active site remain in equilibrium throughout the catalytic cycle, with a switch between the predominate form^[24]^.

In addition to the equilibrium shifts, line shape analysis of the Schiff base nitrogen based on the limiting chemical shifts given by the computational modeling (see the SI) gives direct experimental access to the rates of the tautomeric exchange and hence the activation energy. Interestingly, whereas homogeneous nitrogen linewidths in the absence of exchange (including the βK87 backbone amide) amount to only around 20 Hz, linewidths for the Schiff base nitrogen are on the order of 270 Hz (see Fig. 4C). The Schiff base proton has a linewidth of 190 Hz, compared to amide H^N^ widths of generally around 50 Hz. The exchange-broadened lines suggest a turnover in the μs motional regime with a forward rate of around 5.1 x 10^5^ s^-1^. Even though the solid-state NMR linewidth does not purely reflect the exchange contribution as in solution NMR and would have to be scaled down slightly by an unknown degree, the value obtained of around 41 kJ/mol represents a too large activation barrier for an isolated (de-)protonation event and thus likely reflects the reorganization energy necessary to support the new electrostatic environment. Such remnants of the conformational flexibility between the known crystallographic states would also be in agreement with the slightly elevated *R*_1ρ_ rates in the entry portal for serine in the β-subunit active site as well as the existence of two conformations of the tip of R647 (βR379) in the according X-ray structure. With rates of 5.4 for G352 (βG84) and 8.7 s^-1^ for D649 (βD381), front left in Fig. 3A, often found in mobile residues of otherwise rigid proteins,^[56–58]^ in comparison to rates of only around ~1 s^-1^ generally observed for immobile/inconspicuous residues in this preparation, the slight degree of plasticity in the gate at room temperature is an interesting feature becoming accessible for more in-depth studies. Fig. 4D shows the free-energy landscape of the reaction as derived from these calculations. Actual relaxation dispersion studies, which would be needed to validate/quantify possible exchange contributions to the *R*_1ρ_ rates, are outside the scope of this study due to the exhaustive measurement time associated with multiple pseudo-4D relaxation experiments (i. e., a pseudo-5D) at 700 MHz. Nevertheless, with the bottleneck of obtaining a large share of the TS assignment having been overcome, this study has finally leveraged an access to detailed assessment of the conformational dynamics associated with the allosteric control in TS as well as its connection to Schiff base proton exchange and other specific catalytic properties within the active sites long sought.

## Discussion

The chemical shift in NMR is a sensitive probe of the individual chemical environment of each atom, reporting on protonation state, hybridization state, hydrogen bonding interactions, and the surrounding electrostatic environment. But unlocking this information through the assignment of chemical shifts remains elusive in larger, more complex proteins. Given the difficulty of even employing much simpler sequences like the 3D HNCACB for large globular proteins in solution,^[59,60]^ it is an important realization that solid-state NMR, in contrast to regular solution NMR, can effectively exploit higher-dimensionality approaches for systems of very high molecular weight.

The assignments in TS are representative for a large number of other high-molecular-weight targets in biology, pharmacological research, and biotechnology. For example, common drug targets, including many kinases, phosphatases, nuclear receptors, and many membrane proteins like surface receptors, channels, and transporters, are often in a molecular-weight range of 60 to 80 kDa.^[61–64]^ Similarly, the monomeric molecular mass of many biocatalysts in industrial applications like dehydrogenases, lipases and esterases often fall into this molecular-weight regime.^[65]^ Whereas the general advantages of solid-state NMR for symmetric complexes – as sensitivity is not compromised by homo-oligomerization – or membrane proteins are generally agreed upon, solid-state NMR thus also bears promise for characterizing monomeric proteins or asymmetric complexes beyond what is thought to be possible nowadays. Whereas on the 700 MHz system, an extended amount of measurement time was required for this study yet, much shorter measurement times will be needed for higher fields, further emphasizing the perspectives of the approach.

The SI shows spectra and sensitivities obtained for a non-deuterated TS sample spun at 111 kHz MAS in a 0.7 mm rotor, which would circumvent the likely problem of incomplete back-exchange into the deuterated sample. In fact, very similar experiments are possible as for deuterated samples, as exemplified by a 4D hCOCANH experiment recorded for comparison (Fig. S9). Lower sensitivity due to reduced sample volume (~0.5 μl), however, in combination with the general measurement time requirements for the 72 kDa sample (vide supra), make this a non-trivial problem. With increasing time resources available on very fast spinning probes generally, or labeling schemes that foster high deuteration without back-exchange or stochastic proton occupancy^[66]^, however, we expect near-complete assignments can become straightforward to attain.

Chemical shift measurements in TS have been the preeminent tool for characterizing the chemical structures of intermediates throughout the catalytic cycle, highlighting the protonation states and associated tautomeric equilibria for the cofactor, substrates, and active-site residues.^[39–41,47,67,68]^ To date, this approach has relied heavily on the production of multiple protein samples for solid-state NMR with distinct, selective incorporation of ^13^C and ^15^N spin labels, and the subsequent use of the measured shifts as restraints in first-principles computational refinement. As these *in-silico* approaches hinge on priors from real, experimentally obtained shifts, the new level of comprehensiveness for the chemical shift data obtained through extensive assignments on U-^13^C,^15^N-labeled samples ushers in new possibilities for NMR-assisted crystallography that we expect will allow it to expand beyond the active site to include extended hydrogen bonding and proton transfer pathways, conformational dynamics, and allostery. We anticipate that the analysis pursued for the resting internal aldimine state studied here can be extended in a reasonably straightforward way to other stable intermediates along the catalytic pathway, as additional intermediates would not largely alter the chemical shifts for the majority of the enzyme and the bulk of the information can be transferred.

## Conclusions

Here we have shown the first protein solid-state NMR study, enabled by higher-dimensionality (4D and 5D) shift assignments, in the 144 kDa, bi-enzyme tryptophan synthase complex. The benefits of higher dimensionality required for the 665 residue asymmetric unit, in particular low-ambiguity sequential correlations directly concatenated with residue type data, are enabled in the solid state by high transfer efficiencies irrespective of molecular weight. In combination with state-of-the-art computational approaches, these provide access to chemical, thermodynamic, and kinetic parameters for active site species and give first experimental hints of the plasticity that is thought to be essential for substrate trapping and product release. This study ushers in a new molecular-weight regime for protein solid-state NMR, demonstrating the feasibility of assignment and assessment of dynamics and chemical properties for much larger proteins than previously thought, even with a 700 MHz spectrometer. Opening up this size regime for solid-state NMR and NMR generally will have extensive implications for access to an atomic-level understanding of reaction thermodynamics and kinetics widely sought for biological, medical, and industrial applications.

## Acknowledgements

We thank the group of W. Koźmiński and Jan Stanek for helpful discussions about SSA and related processing scripts. Financial support is acknowledged from the Center of NanoScience (CeNS) and the Deutsche Forschungsgemeinschaft (DFG, German Research Foundation) in the context of SFB 749, TP A13 (project number 27112786), SFB 1309, TP 03 (project number 325871075), and the Emmy Noether program. This work was funded under Germany’s Excellence Strategy – EXC-2033 and EXC-114 – project numbers 390677874 and 24286268 – and United States’ NSF Grant CHE1710671 and NIH Grants GM097569 and GM137008 given to L.J.M. Gefördert durch die Deutsche Forschungsgemeinschaft (DFG) – SFB 1309 – 325871075; SFB 749 – 27112786. Gefördert durch die Deutsche Forschungsgemeinschaft (DFG) im Rahmen der Exzellenzstrategie des Bundes und der Länder – EXC 2033 – Projektnummer 390677874; EXC 114–24286268. The authors gratefully acknowledge the computing time provided on the Linux HPC cluster at Technical University Dortmund (LiDO3), partially funded in the course of the Large-Scale Equipment Initiative by the German Research Foundation (DFG) as project 271512359. Additional computations were performed using the computer clusters and data storage resources of the UC Riverside HPCC, which were funded by grants from United States NSF (MRI-1429826) and NIH (S10OD016290).

